## Supporting Information for "TopEC: Improved classification of enzyme function by a localized 3D protein descriptor and 3D Graph Neural Networks"

#### Supplemental Data

**Data S1.** PyCM reports for all networks trained for the manuscript.

**Data S2.** Modified Price and PrOSPECCTs dataset. Contains a list of PDBs for each sub dataset.

### Supporting Tables

**Table S1.** The pair-wise sequence identity for structures in Fig. 4g.

| **PDB** | 1QF6 | 3UH0 | 3UGQ | 5ZY9 | 6VU9 |
|---|---|---|---|---|---|
| 1QF6 | 100% | | | | |
| 3UH0 | 38% | 100% | | | |
| 3UGQ | 38% | 100% | 100% | | |
| 5ZY9 | 40% | 43% | 43% | 100% | |
| 6VU9 | 59% | 37% | 36% | 40% | 100% |

**Table S2.** Atom annotation per residue type.

| **Backbone Atoms** | | |
|---|---|---|
| *13 in GLY/PRO<br>**Not present in GLY/PRO | N | 1 |
| | C | 2 |
| | CA* | 3 (13*) |
| | O / OXT | 4 |
| | H** | 32 |
| | HA** | 33 |

| **Neutral-Nonpolar R Groups** | | |
|---|---|---|
| Glycine **GLY** | H | 32 |
| | 1HA / 2HA | 39 |
| Alanine **ALA** | CB | 8 |
| | 1HB / 2HB / 3HB | 36 |
| Valine **VAL** | CB | 7 |
| | CG1 / CG2 | 8 |
| | HB | 34 |
| | 1HG1 / 2HG1 / 3HG1 / 1HG2 / 2HG2 / 3HG2 | 36 |
| Isoleucine **ILE** | CB | 6 |
| | CG1 | 7 |
| | CG2 / CD1 | 8 |
| | HB | 34 |
| | 1HD1 / 2HD1 / 3HD1 / 1HG2 / 2HG2 / 3HG2 | 36 |
| | 1HG1 / 2HG1 | 35 |
| Leucine **LEU** | CB | 7 |
| | CG | 6 |
| | CD1 / CD2 | 8 |
| | HG | 34 |
| | 1HB / 2HB | 35 |

| | | |
|---|---|---|
| | 1HD1 / 2HD1 / 3HD1 / 1HD2 / 2HD2 / 3HD2 | 36 |
| Proline **PRO** | CB / CG / CD | 14 |
| | 1HB / 2HB / 1HD / 2HD / 1HG / 2HG | 35 |
| Methionine **MET** | CB / CG | 7 |
| | CE | 8 |
| | SD | 22 |
| | 1HB / 2HB / 1HG / 2HG | 35 |
| | 1H / 2H / 3H / 1HE / 2HE / 3HE | 36 |

| **Neutral-Polar R Groups** | | |
|---|---|---|
| Serine **SER** | CB | 7 |
| | OG | 10 |
| | 1HB / 2 HB | 35 |
| | HG | 38 |
| Threonine **THR** | CB | 6 |
| | CG2 | 8 |
| | OG1 | 10 |
| | HB | 34 |
| | 1HG2 / 2HG2 / 3HG2 | 36 |
| | HG | 38 |
| Asparagine **ASN** | CB | 7 |
| | CG | 5 |
| | ND2 | 17 |
| | OD1 | 12 |
| | 1HB / 2HB | 35 |
| | 1HD2 / 2HD2 | 43 |
| Glutamine **GLN** | CB / CG | 7 |
| | CD | 5 |
| | NE2 | 17 |
| | OE1 | 12 |
| | 1HB / 2HB / 1HG / 2HG | 35 |
| | 1HE2 / 2HE2 | 43 |
| Cysteine **CYS** | CB | 7 |
| | SG | 20 |
| | 1HB / 2HB | 35 |
| | HG | 45 |

| Acidic R Groups | | |
|---|---|---|
| Aspartic Acid **ASP** | CB | 7 |
| | CG | 5 |
| | OD1 / OD2 | 11 |
| | 1HB / 2HB | 35 |
| Glutamic Acid **GLU** | CB / CG | 7 |
| | CD | 5 |
| | OE1 / OE2 | 11 |

| Basic R Groups | | |
|---|---|---|
| Lysine **LYS** | CB / CG / CD / CE | 7 |
| | NZ | 26 |
| | 1 HB / 2HB / 1HD / 2 HD / 1 HE / 2HE / 1HG / 2HG | 35 |
| | 1HZ / 2HZ / 3HZ | 47 |
| Arginine **ARG** | CB / CG / CD | 7 |
| | CZ | 5 |
| | NE | 15 |
| | NH1 / NH2 | 16 |
| | HE | 41 |
| | 1HB / 2HB / 1HD / 2HD / 1HG / 2HG | 35 |
| | 1HH1 / 2HH1 / 1HH2 / 2HH2 | 42 |

| Aromatic R Groups | | |
|---|---|---|
| Histidine **HIS** | CB | 7 |
| | CG | 28 |
| | CD | 27 |
| | CE1 | 25 |
| | ND1 | 15 |
| | NE2 | 15 |
| | 1HB / 2HB | 35 |
| | HD1 / HE2 | 41 |
| | HE1 | 46 |
| | HD2 | 48 |
| Phenylalanine **PHE** | CB | 7 |
| | CG | 21 |
| | CD1 / CD2 / CE1 / CE2 / CZ | 9 |
| | HD1 / HD2 / HE1 / HE2 / HZ | 37 |

| | | |
|---|---|---|
| | 1HB / 2HB | 35 |
| Tryptophan **TRP** | CB | 7 |
| | CG | 29 |
| | CD1 / CE3 / CZ3 | 9 |
| | CD2 / CE2 | 24 |
| | CH2 / CZ2 | 18 |
| | NE1 | 15 |
| | 1HB / 2HB | 35 |
| | HD1 / HE3 / HZ3 | 37 |
| | HE1 | 41 |
| | HH2 / HZ2 | 44 |
| Tyrosine **TYR** | CB | 7 |
| | CG | 23 |
| | CD1 / CD2 / CE1 / CE2 | 9 |
| | CZ | 19 |
| | OH | 10 |
| | HD1 / HD2 / HE1 / HE2 | 37 |
| | HH | 38 |
| | 1HB / 2HB | 35 |

### Supporting Figures

**Supporting Figure 1.** Model performance when training with and without oversampling.

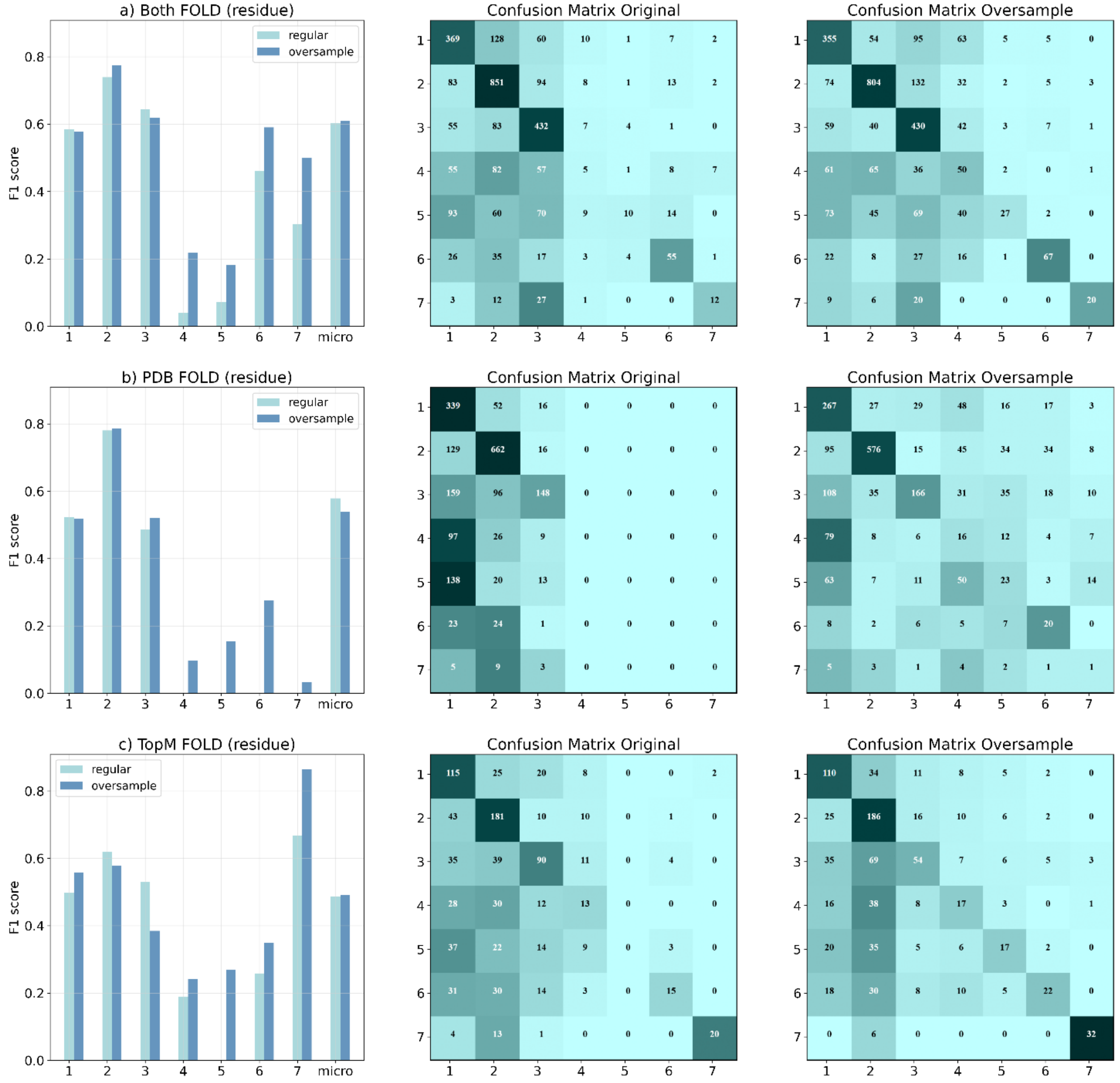

Left: The F-score for each mainclass of the fold networks tested in Table 1A. Middle: The confusion matrix for the networks tested without oversampling. Right: The confusion matrix for the networks tested with oversampling.

**Supporting Figure 2.** EnzyNet results for all modes and datasets tested.

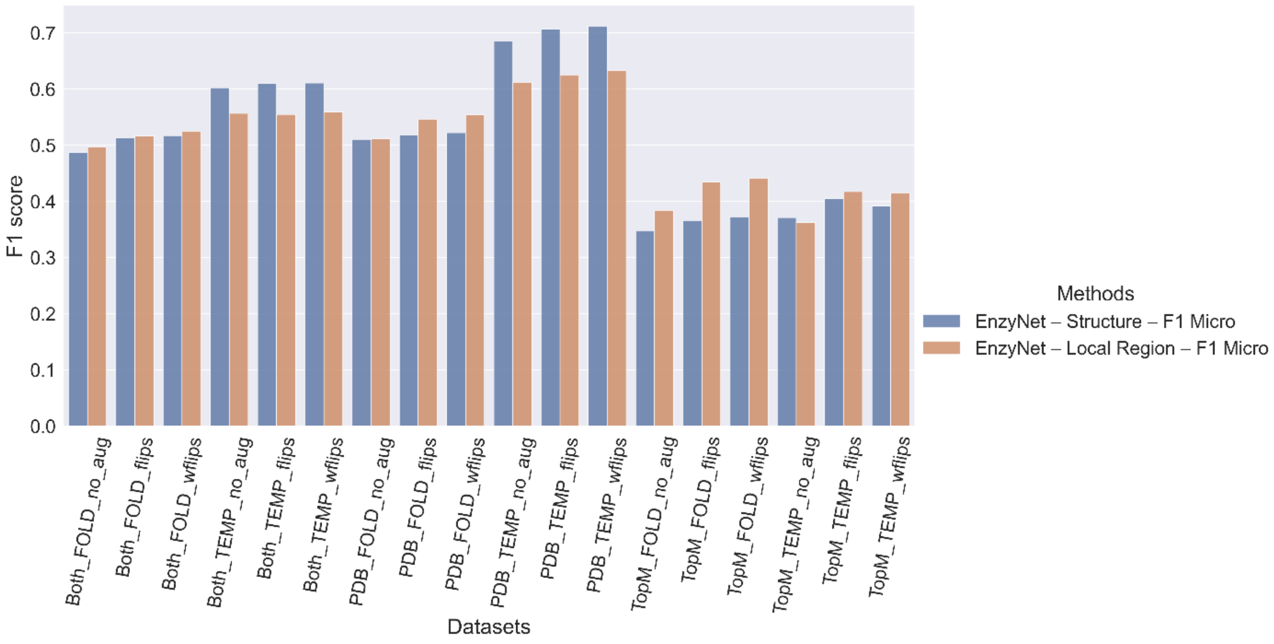

Each dataset and split for networks trained on the full structure and the local region (16 Å around the binding site). For each dataset and split combination, we tested three methods: no augmentation (*_no_aug),* flips (*_flips),* and weighted flips *(_wflips)*.

**Supporting Figure S3.** F1 score as a function of graph node count for hierarchical EC classification.

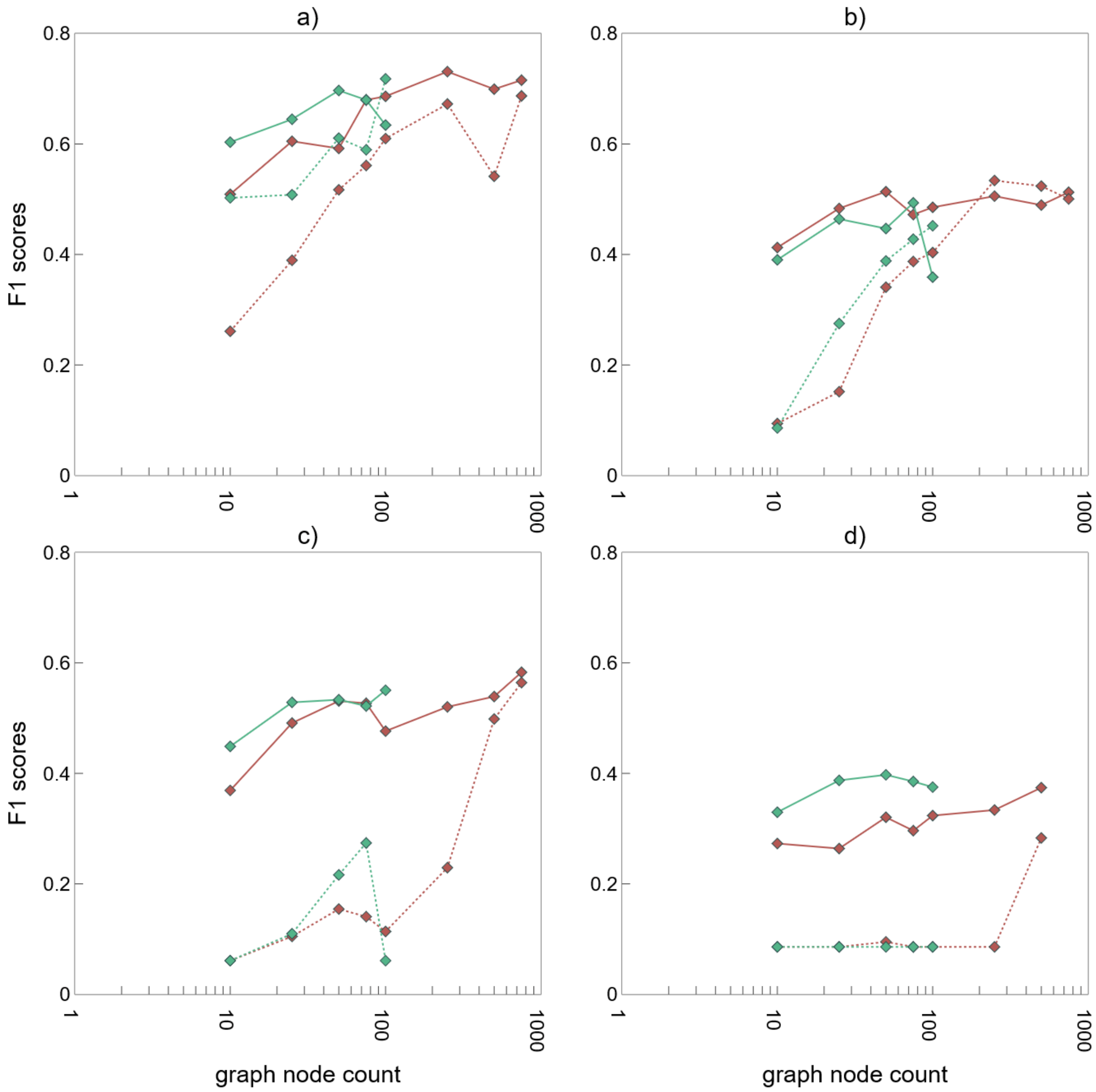

Red: Networks trained with a SchNet model. Green: Networks trained with a DimeNet++ model. For the continuous lines, we created the localized 3D descriptors using binding site information. For the dotted lines, we created the localized 3D descriptor from a random point on the protein. We show the result for the combined dataset using a) residue resolution with a temporal split, b) residue resolution with a fold split, c) atom resolution with a temporal split, d) atom resolution with a fold split.

**Supporting Figure S4.** Model performance when training on *ab initio* predicted structures (AlphaFold2) or homology modeled structures (TopModel) for various descriptors and networks tested.

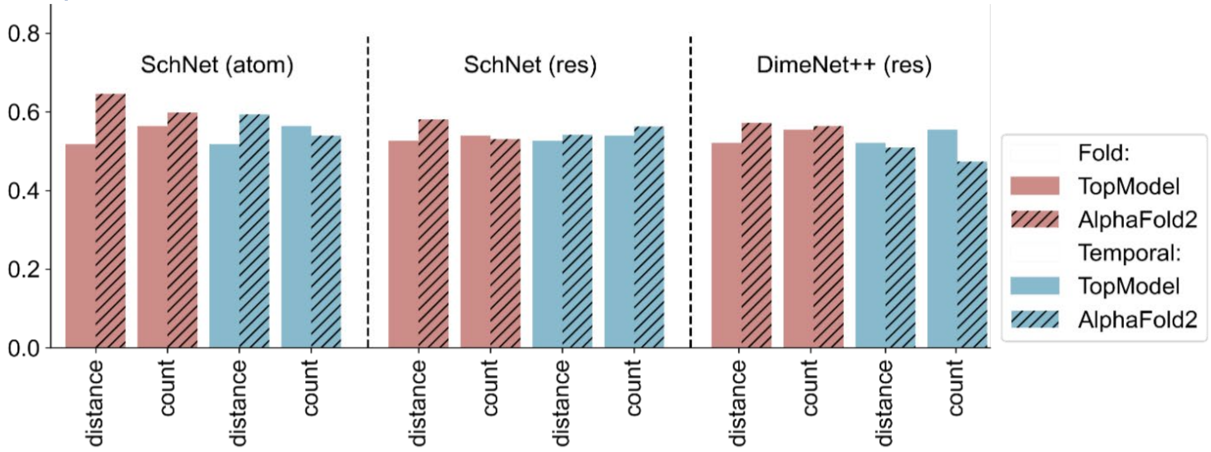

F-score for the networks trained with models obtained from TopModel (no stripes) and AlphaFold2 (stripes) for a fold split (red) and a temporal split (blue).

**Supporting Figure S5.** 10 different folds for type II site-specific deoxyribonucleases.

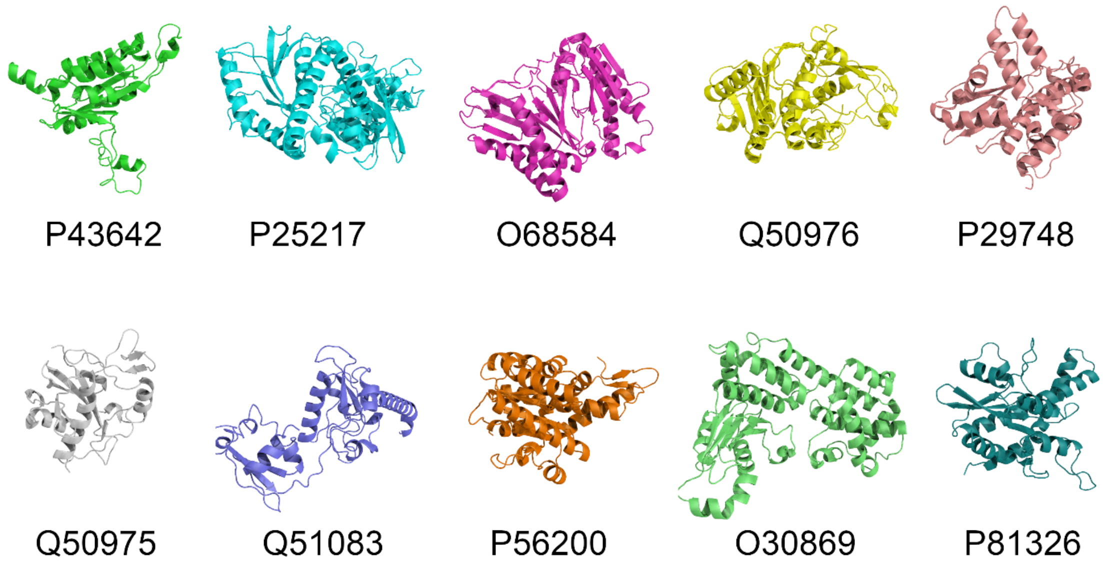

None of the structures are correctly predicted by a network trained on AF703 using a random split.

**Supporting Figure S6.** The area under the precision-recall curve against different properties of the data.

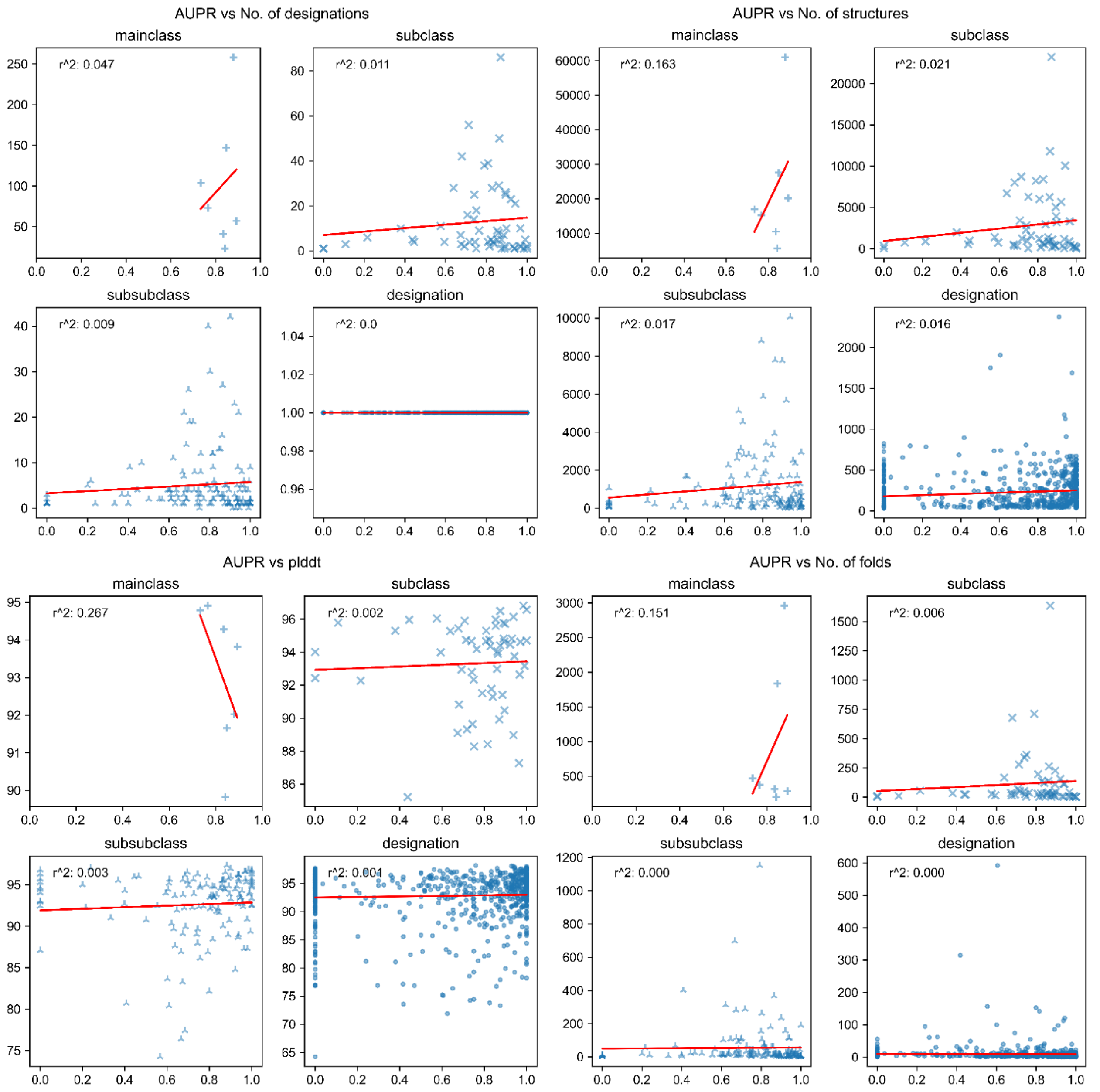

**Supporting Figure S7.** F1 score for all Price and ProSPECCTs datasets broken down by tested network type.

| | Price | DS1 | DS1.2 | DS2 | DS3 | DS4 | DS5 | DS5.2 | DS6 | DS6.2 | DS7 |
|---|---|---|---|---|---|---|---|---|---|---|---|
| **fold_SchNet_atom_distance** | 0.24 | 0.12 | 0.44 | 0.48 | 0.37 | 0.36 | 0.48 | 0.49 | 0.69 | 0.69 | 0.55 |
| **fold_SchNet_atom_count** | 0.10 | 0.04 | 0.11 | 0.20 | 0.33 | 0.34 | 0.40 | 0.46 | 0.38 | 0.38 | 0.35 |
| **fold_SchNet_residue_distance** | 0.14 | 0.37 | 0.56 | 0.44 | 0.56 | 0.56 | 0.68 | 0.65 | 0.69 | 0.69 | 0.66 |
| **fold_SchNet_residue_count** | 0.19 | 0.36 | 0.56 | 0.44 | 0.56 | 0.56 | 0.76 | 0.68 | 0.72 | 0.72 | 0.69 |
| **fold_DimeNet++_atom_distance** | 0.19 | 0.22 | 0.22 | 0.43 | 0.36 | 0.36 | 0.44 | 0.43 | 0.59 | 0.59 | 0.53 |
| **fold_DimeNet++_atom_count** | 0.10 | 0.09 | 0.11 | 0.19 | 0.35 | 0.33 | 0.20 | 0.24 | 0.34 | 0.34 | 0.35 |
| **fold_DimeNet++_residue_distance** | 0.10 | 0.38 | 0.56 | 0.53 | 0.22 | 0.30 | 0.48 | 0.51 | 0.55 | 0.55 | 0.64 |
| **fold_DimeNet++_residue_count** | 0.19 | 0.36 | 0.56 | 0.78 | 0.50 | 0.50 | 0.72 | 0.59 | 0.62 | 0.62 | 0.64 |

In the left column, the split, network, resolution, and localized 3D descriptor type for each subset of the Price and ProSPECCTs dataset are given.

**Supporting Figure S8.** Importance gain per binding site and catalytic residue compared to regular residues.

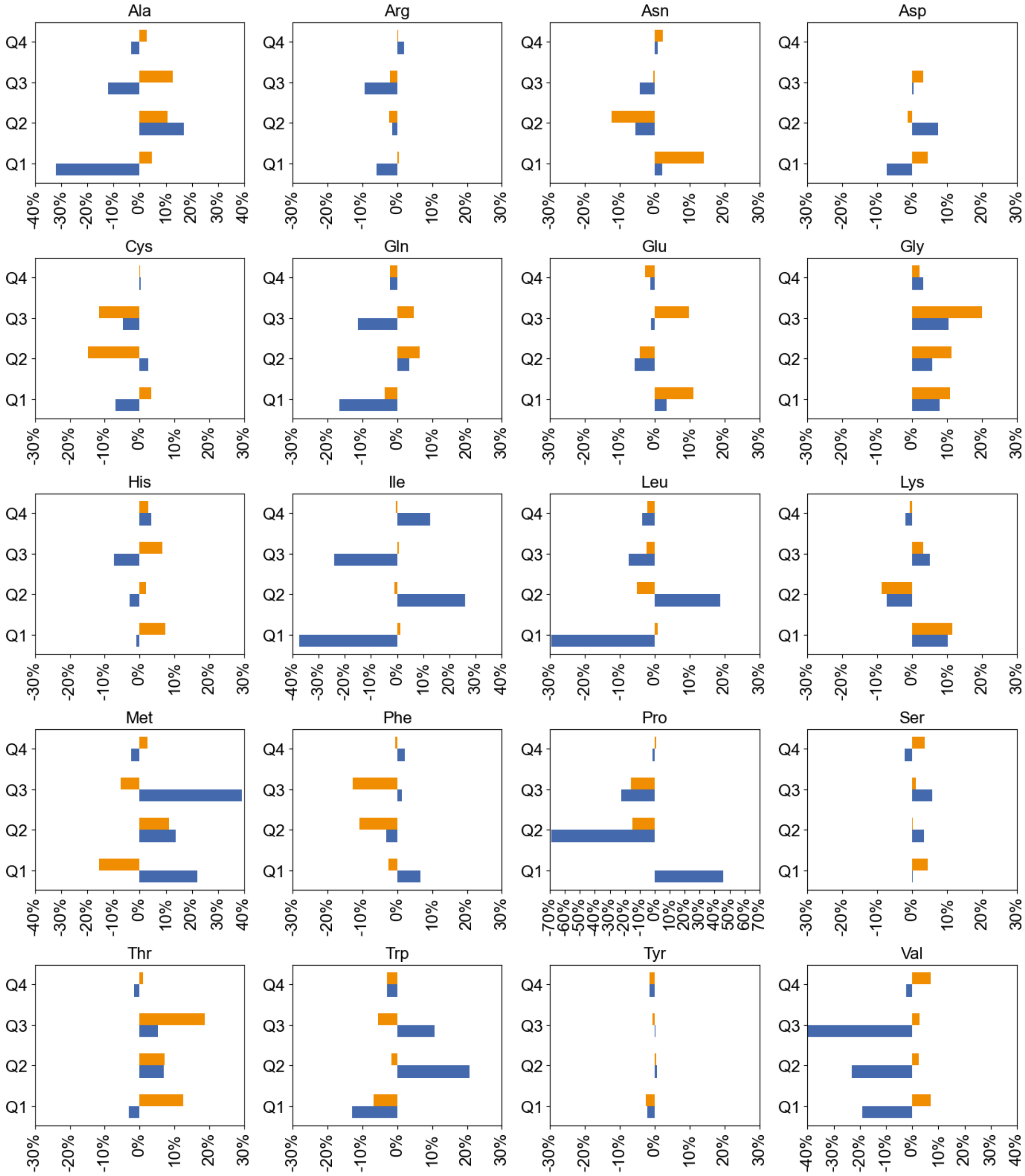

Each panel depicts a separate amino acid. We calculated the importance gain for catalytic (orange) and binding (blue) residues compared to all other residues of that type in the correctly predicted subset of the CSA. We have binned the importance values in four quartiles. A gain in Q1 and Q2 is characterized as positive if we find fewer binding or catalytic residues, while a gain in Q3 and Q4 is characterized as positive if we find more binding or catalytic residues.

**Supporting Figure S9.** Explained PDB structure 1GLA compared to stability predictors.

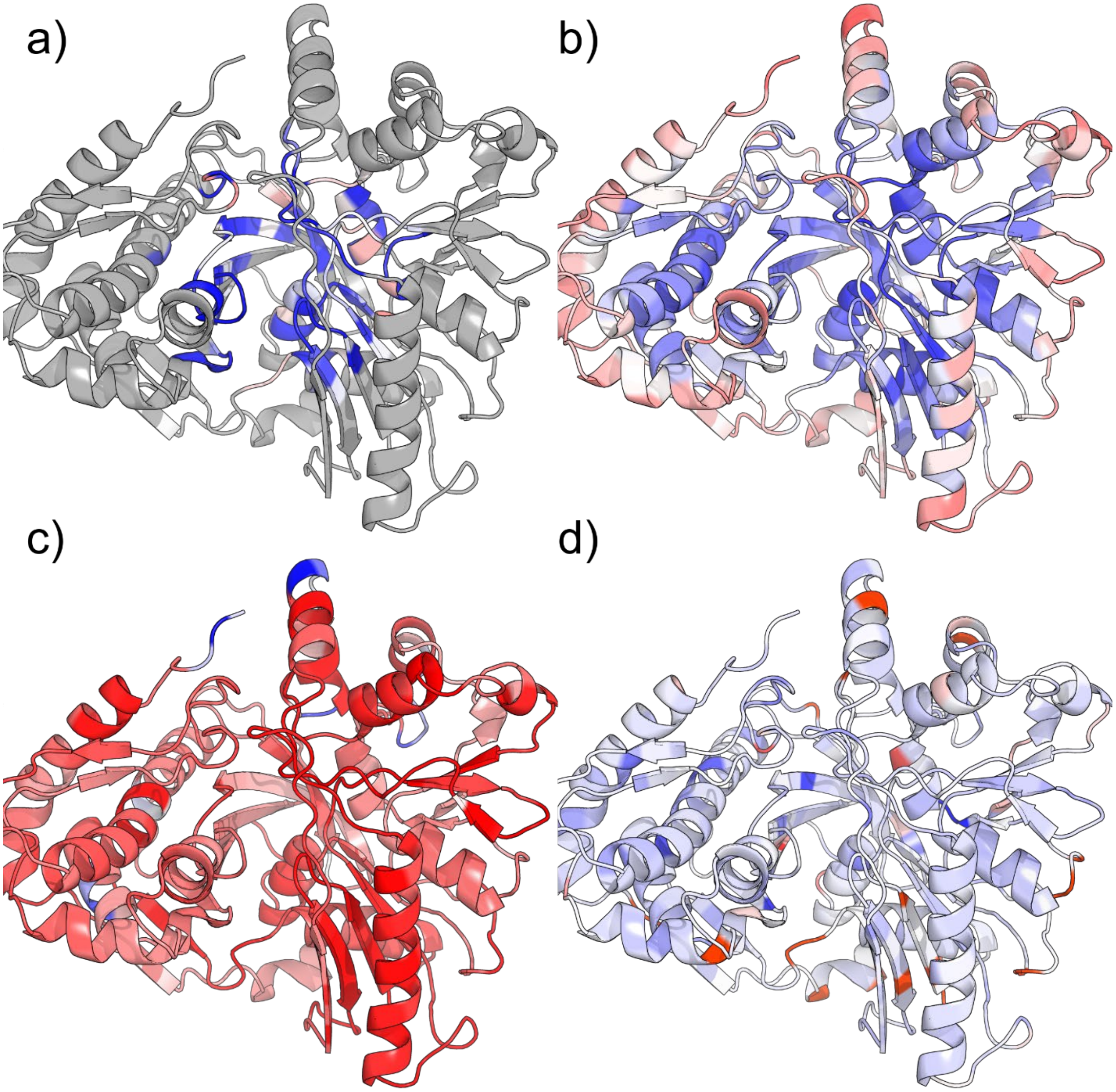

a) GNNExplainer, b) Amino Acids Interaction Webserver, c) Constraint Network Analysis, d) K-Fold. For this structure, we were not able to obtain Thermometer results. Blue indicates stable, red indicates unstable residues.

**Supporting Figure S10.** Explained PDB structure 1WQ1 compared to stability predictors

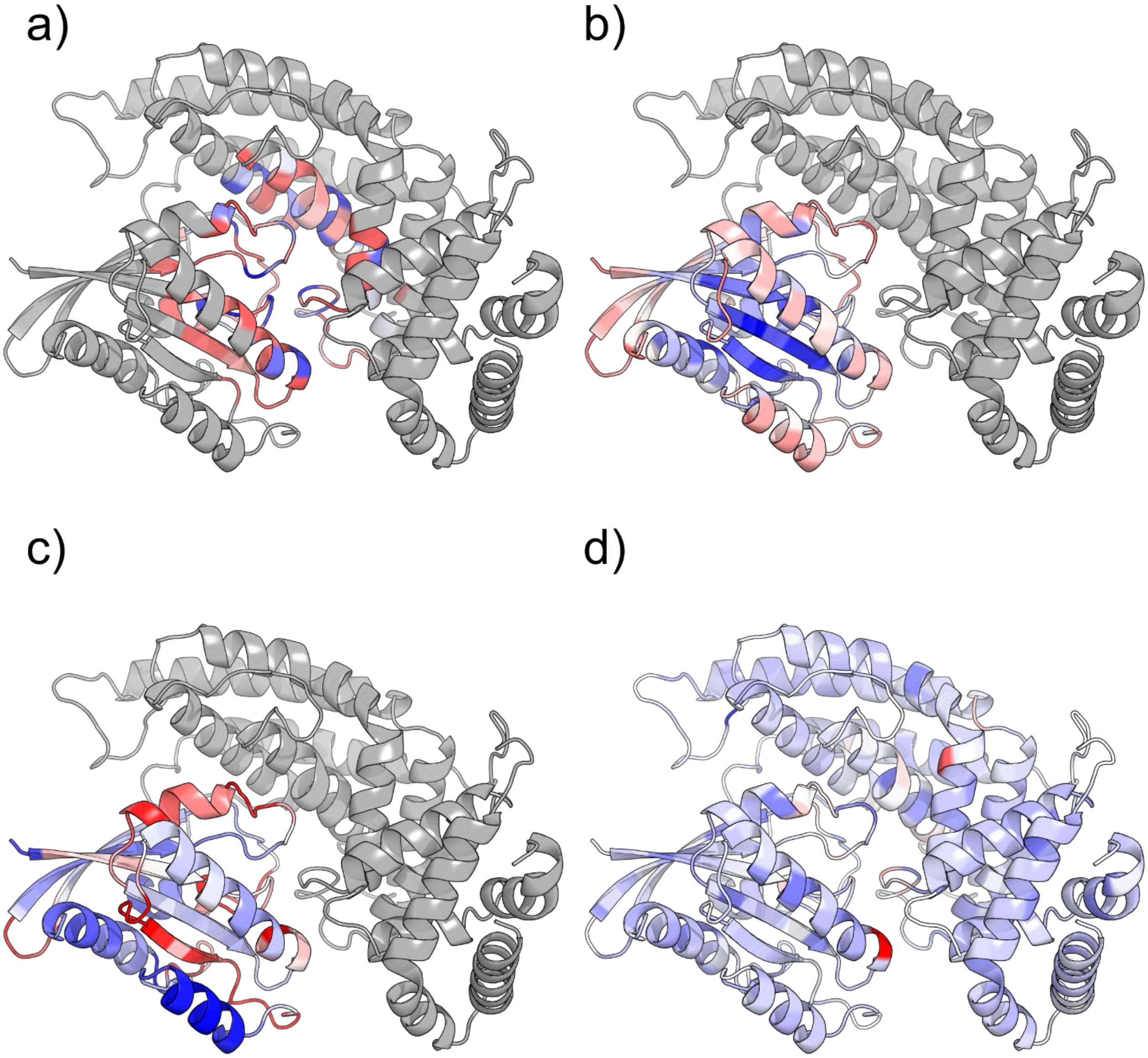

a) GNNExplainer, b) Amino Acids Interaction Webserver, c) Constraint Network Analysis, d) K-Fold. For this structure, we were not able to obtain Thermometer results. Blue indicates stable, red indicates unstable residues.

**Supporting Figure S11.** Explained PDB structure 3BLM compared to stability predictors.

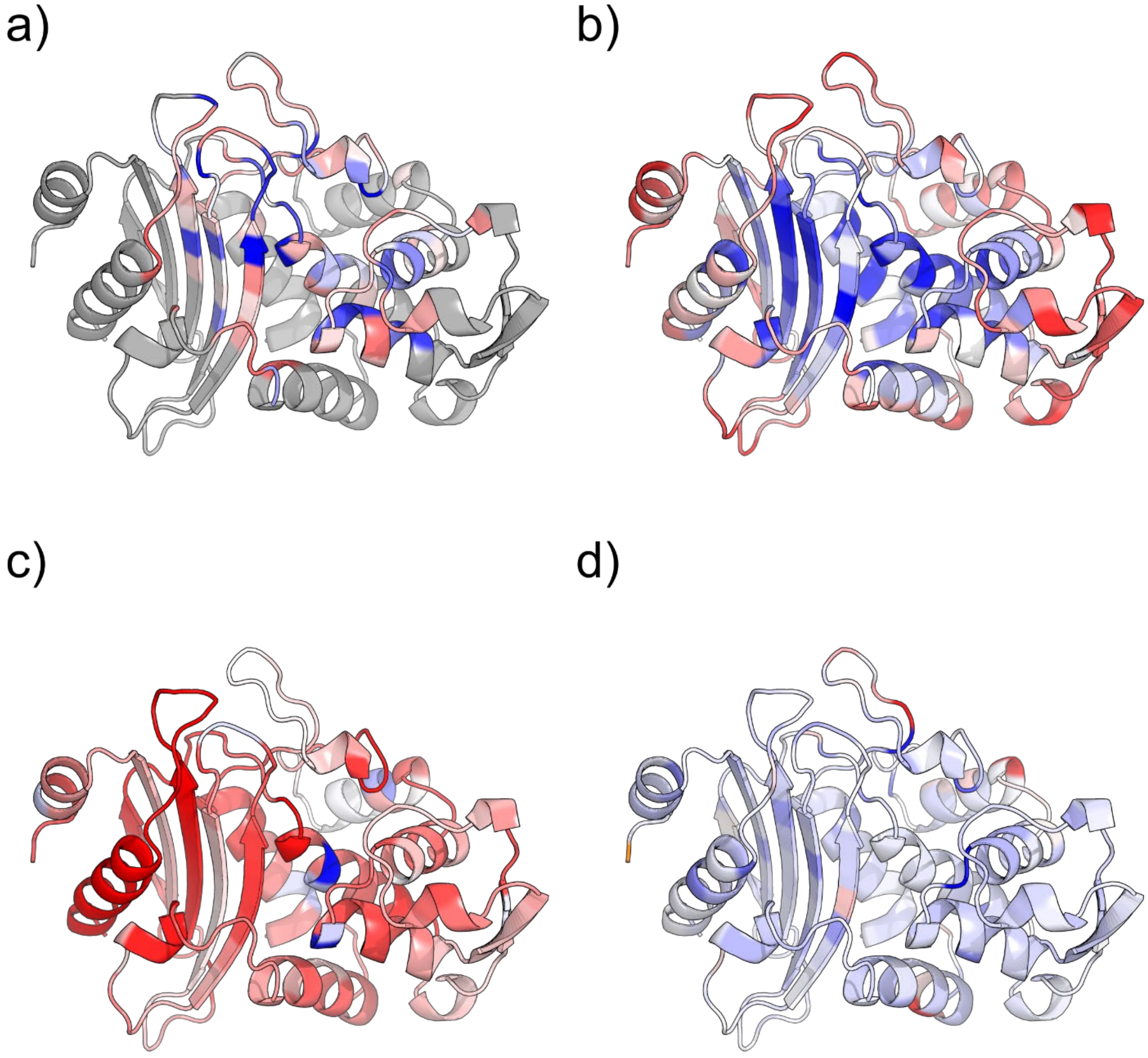

a) GNNExplainer, b) Amino Acids Interaction Webserver, c) Constraint Network Analysis, d) K-Fold. For this structure, we were not able to obtain Thermometer results. Blue indicates stable, red indicates unstable residues.

**Supporting Figure S12.** Explained PDB structure 2MAT compared to stability.

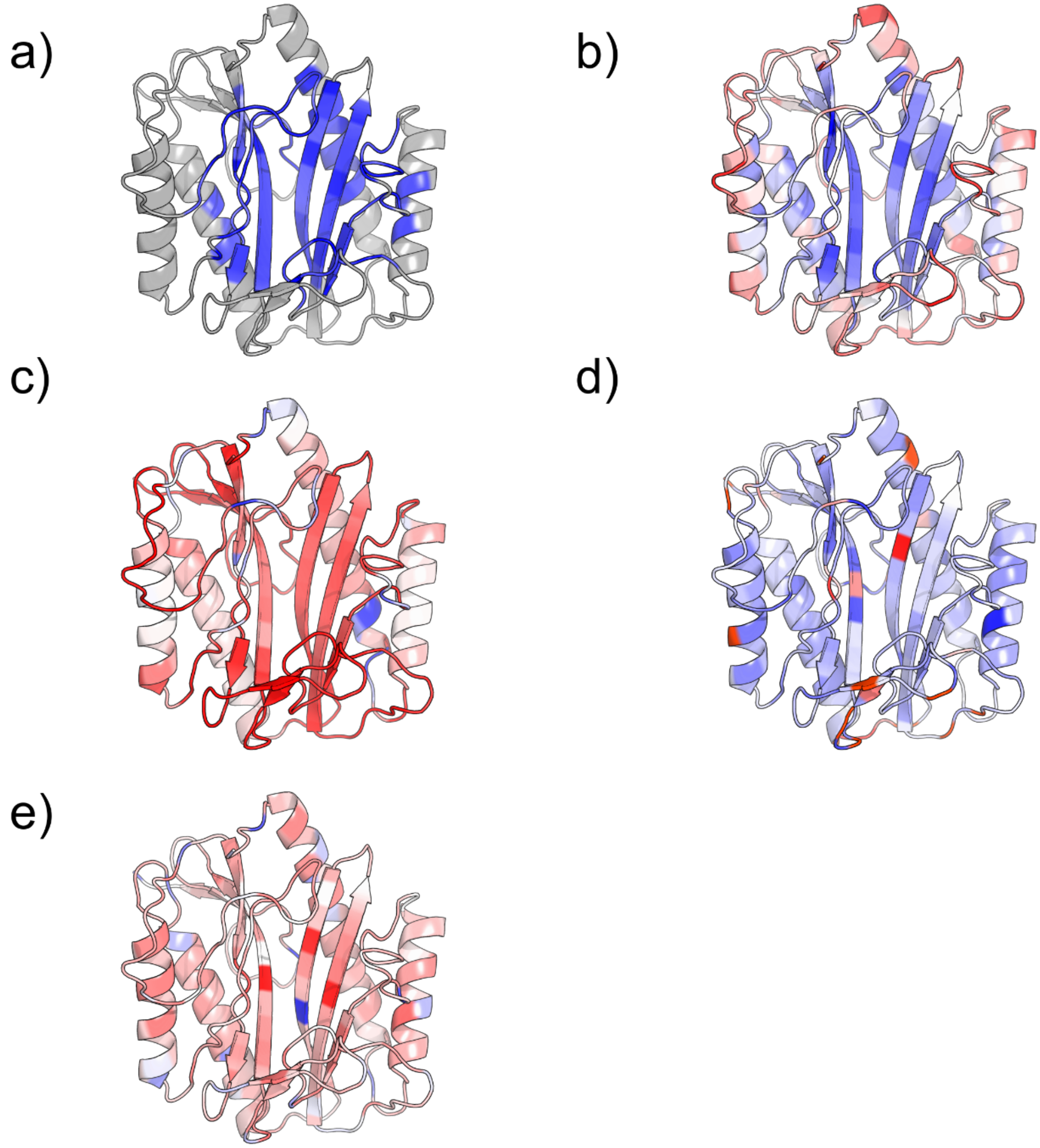

a) GNNExplainer, b) Amino Acids Interaction Webserver, c) Constraint Network Analysis, d) K-Fold, and e) Thermometer. Blue indicates stable, red indicates unstable residues.

**Supporting Figure S13.** Explained PDB structure 1A9U compared to stability predictors.

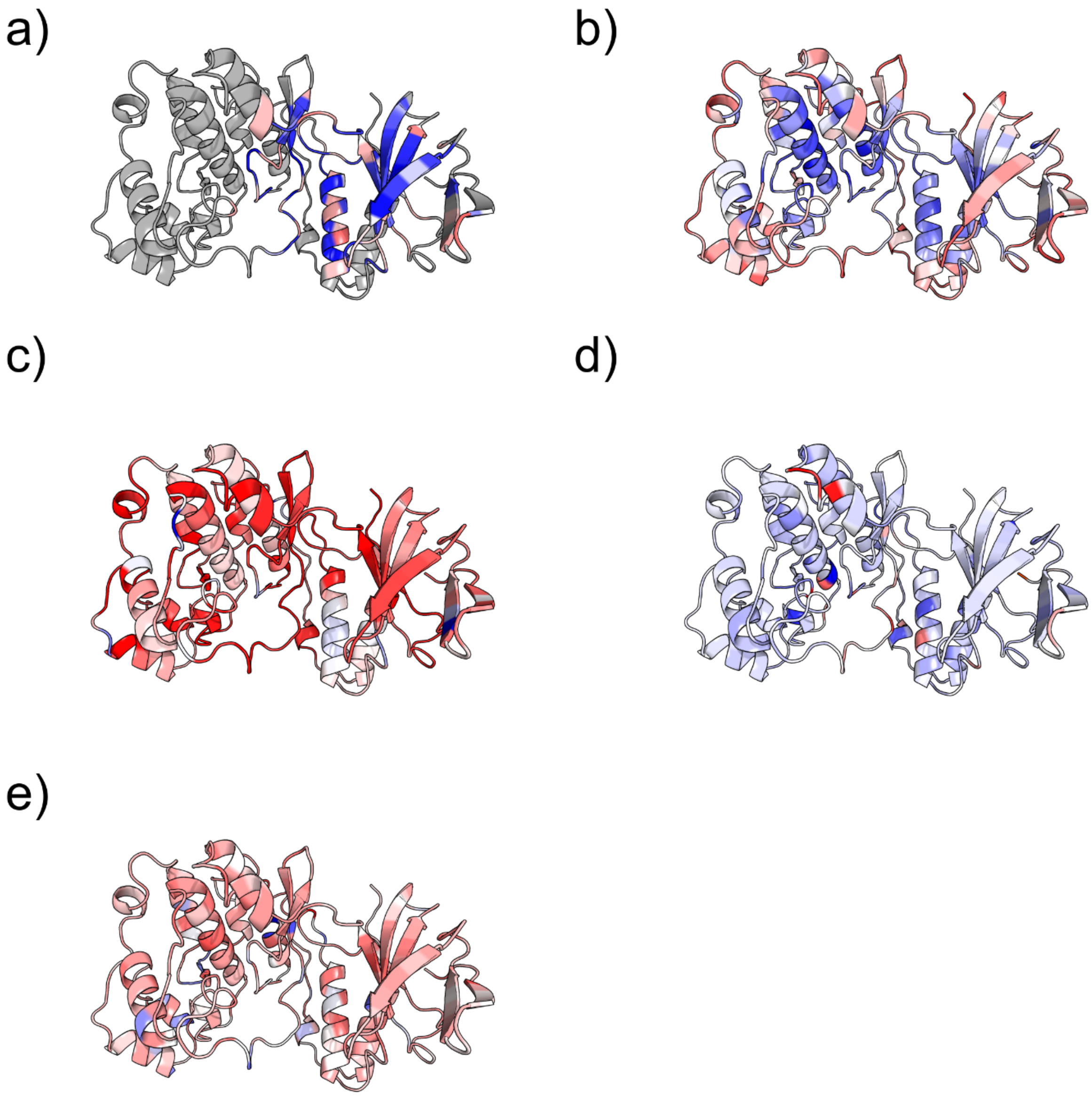

a) GNNExplainer, b) Amino Acids Interaction Webserver, c) Constraint Network Analysis, d) K-Fold, and e) Thermometer. Blue indicates stable, red indicates unstable residues.

**Supporting Figure S14.** Explained PDB structure 3DRC compared to stability predictors.

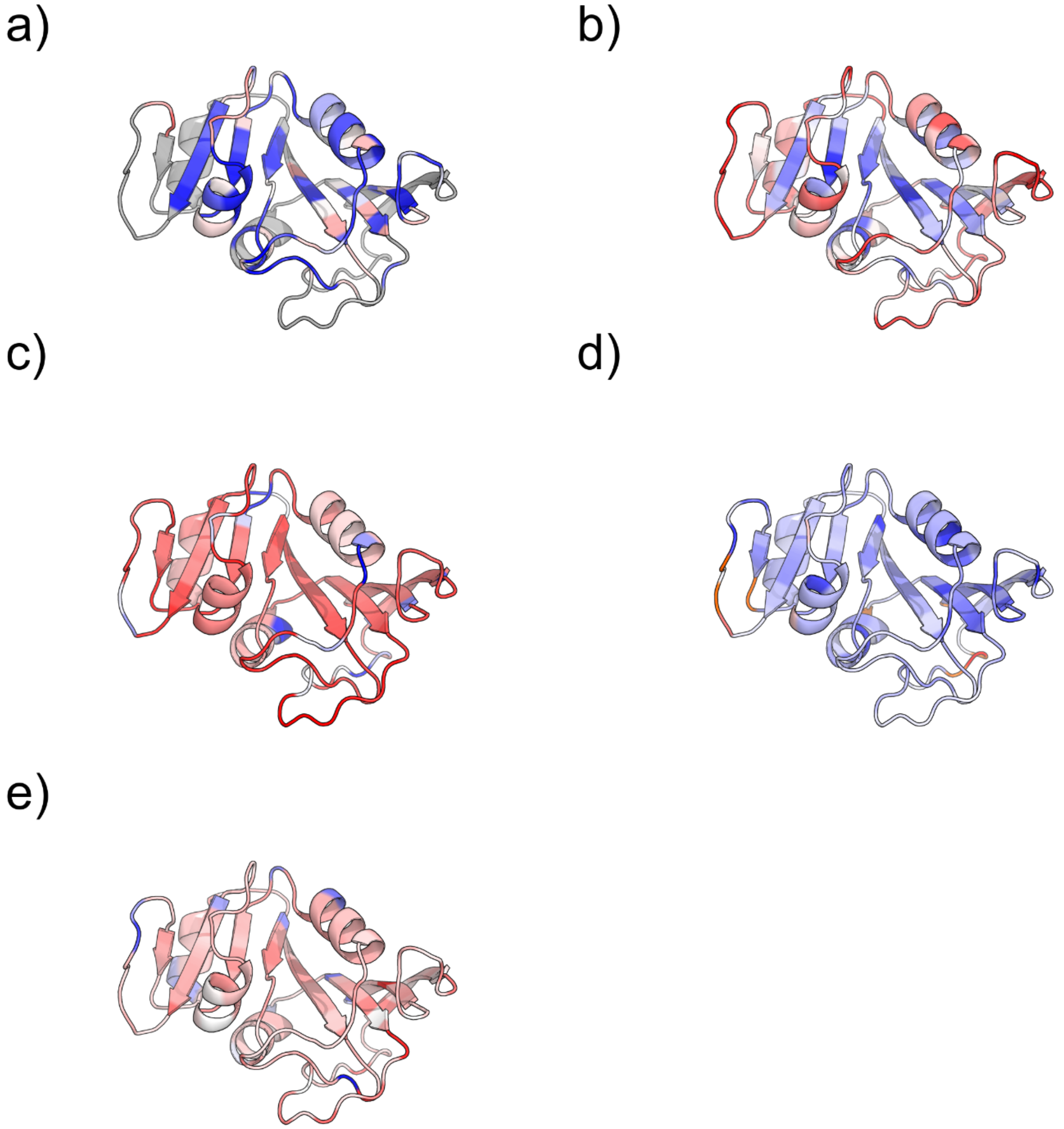

a) GNNExplainer, b) Amino Acids Interaction Webserver, c) Constraint Network Analysis, d) K-Fold, and e) Thermometer. Blue indicates stable, red indicates unstable residues.

**Supporting Figure S15.** Explained PDB structure 1BIW compared to stability predictors.

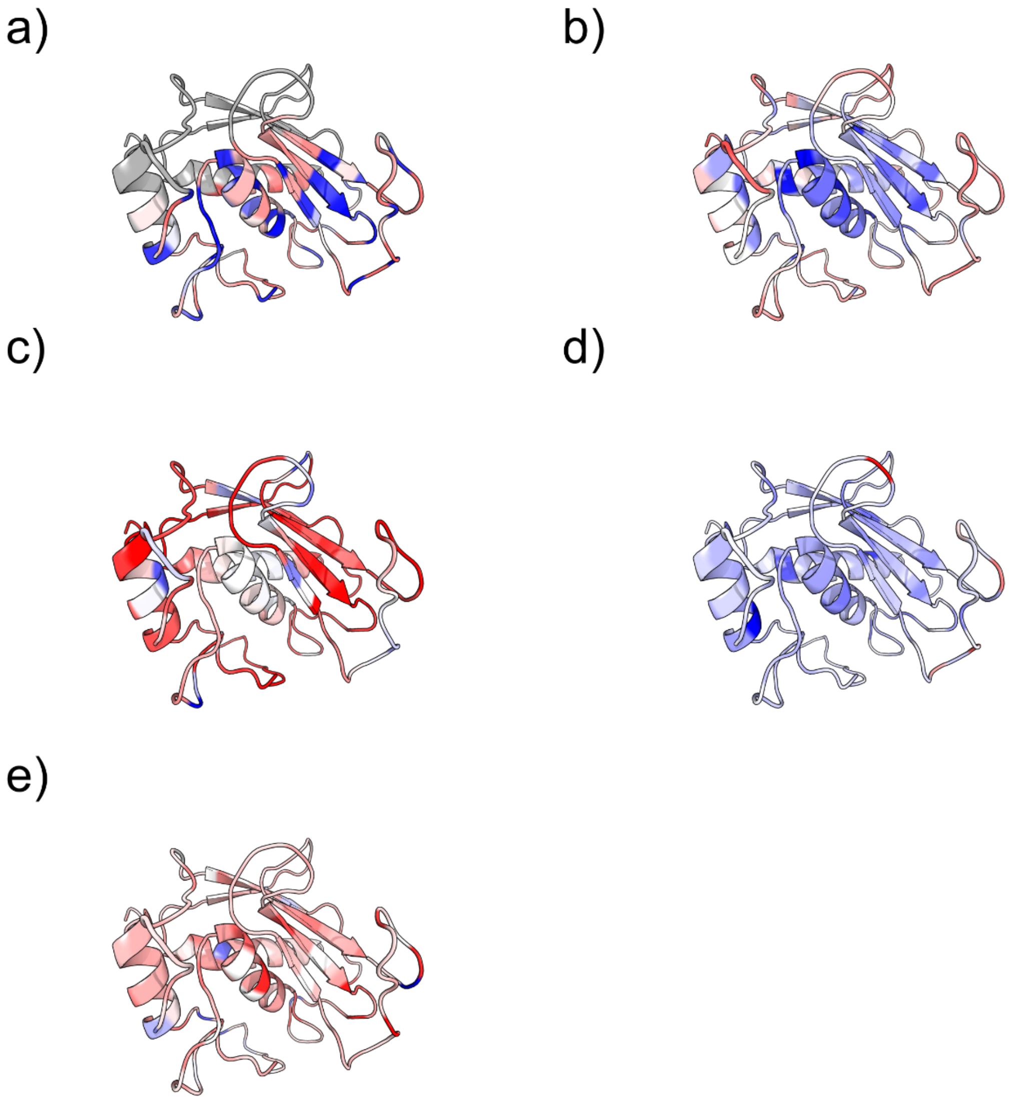

a) GNNExplainer, b) Amino Acids Interaction Webserver, c) Constraint Network Analysis, d) K-Fold, and e) Thermometer. Blue indicates stable, red indicates unstable residues.

**Supporting Figure S16.** Importance for all catalytic and binding atoms.

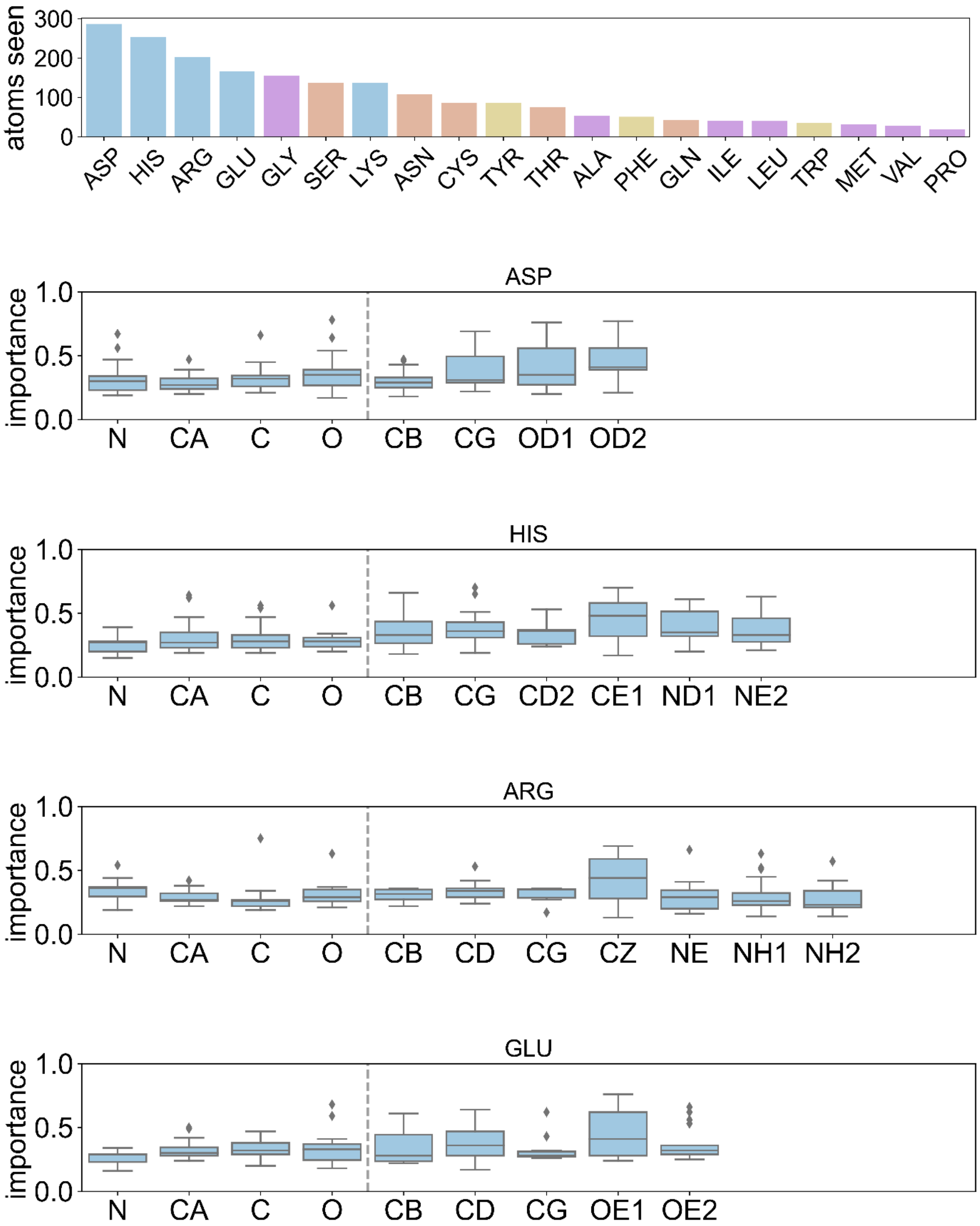

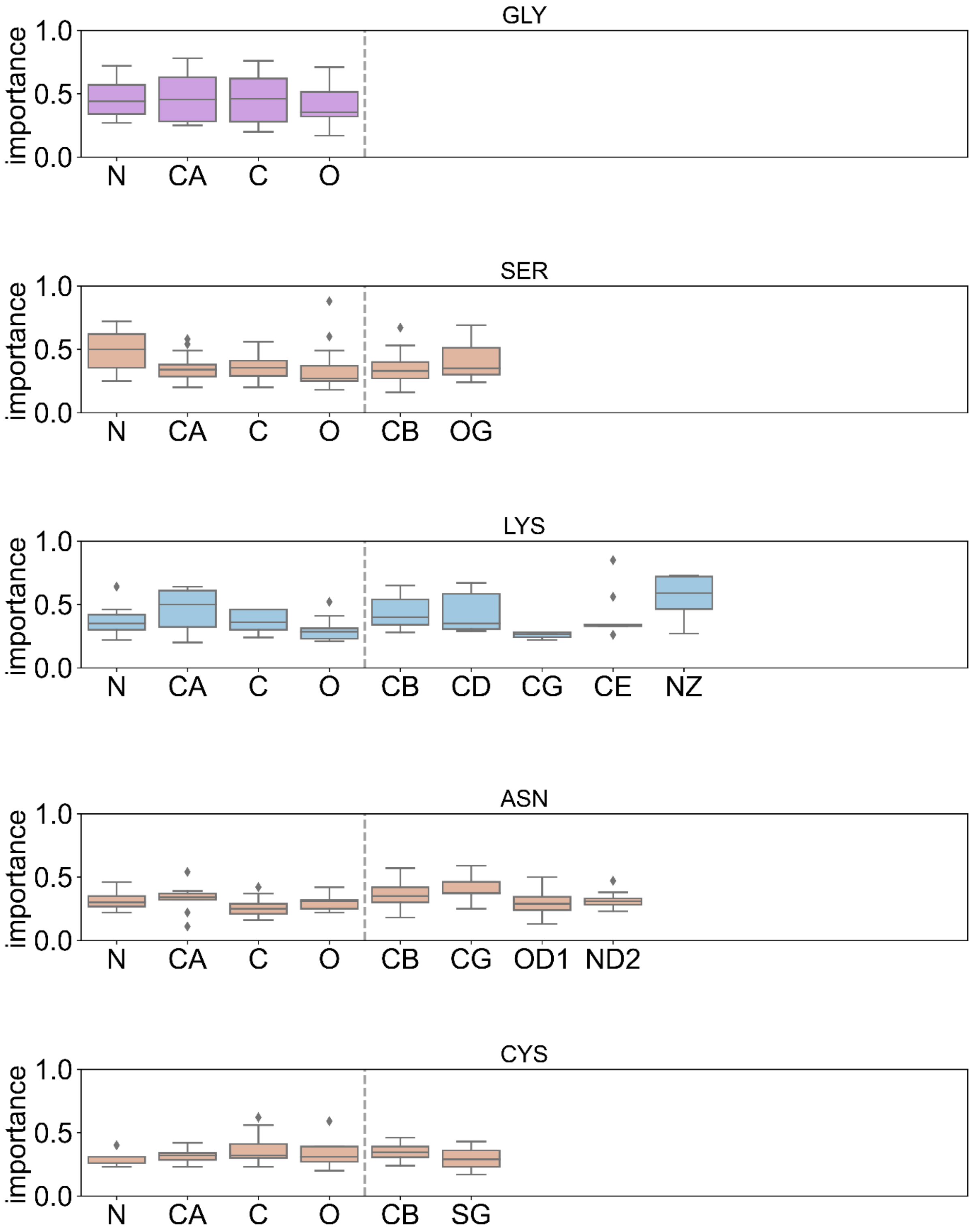
GLY
SER
LYS
ASN
CYS
importance
1.0
0.5
0.0
N
CA
C
O
CB
OG
CD
CG
CE
NZ
OD1
ND2
SG

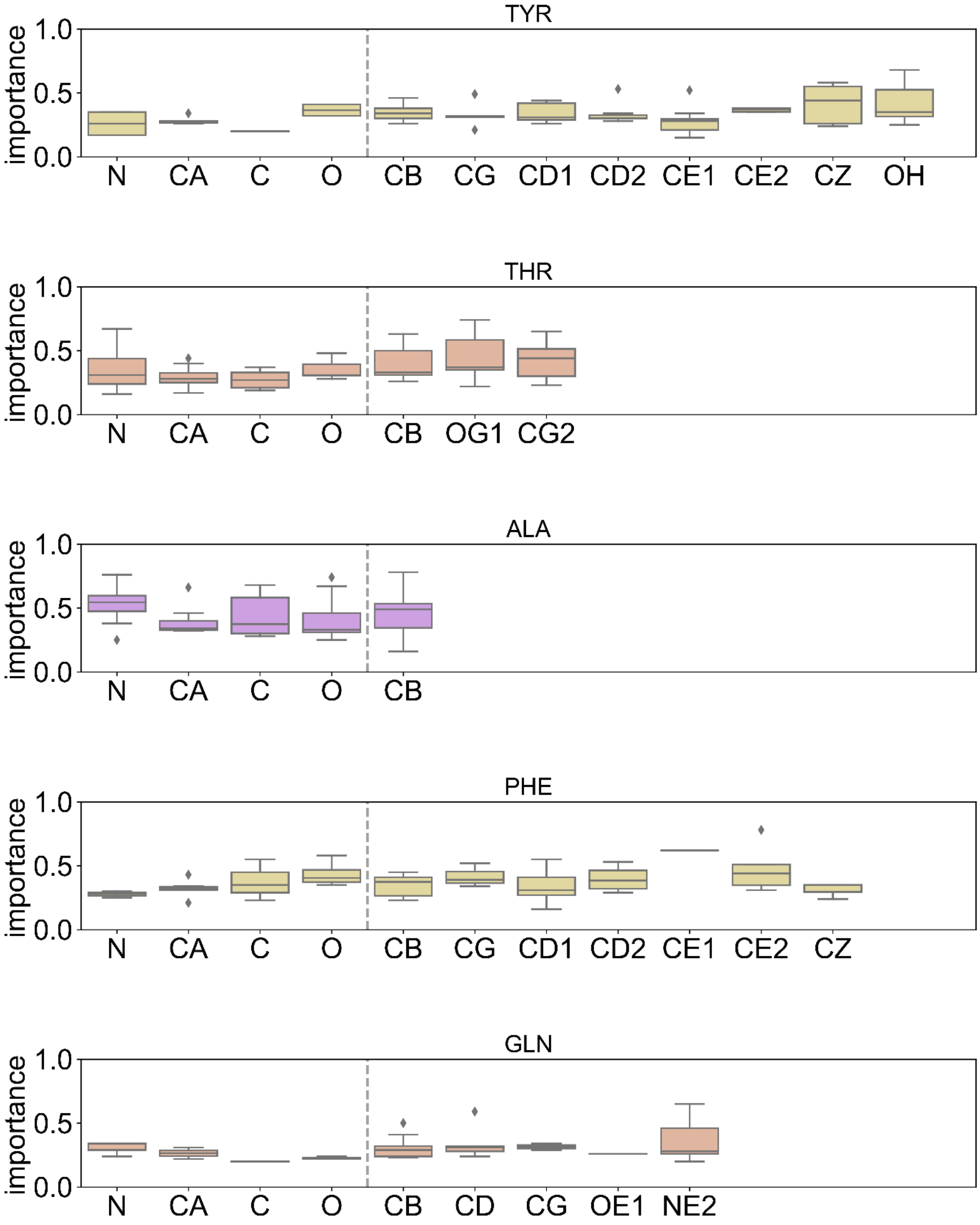
TYR
THR
ALA
PHE
GLN
importance
1.0
0.5
0.0
N CA C O CB CG CD1 CD2 CE1 CE2 CZ OH
N CA C O CB OG1 CG2
N CA C O CB
N CA C O CB CG CD1 CD2 CE1 CE2 CZ
N CA C O CB CD CG OE1 NE2

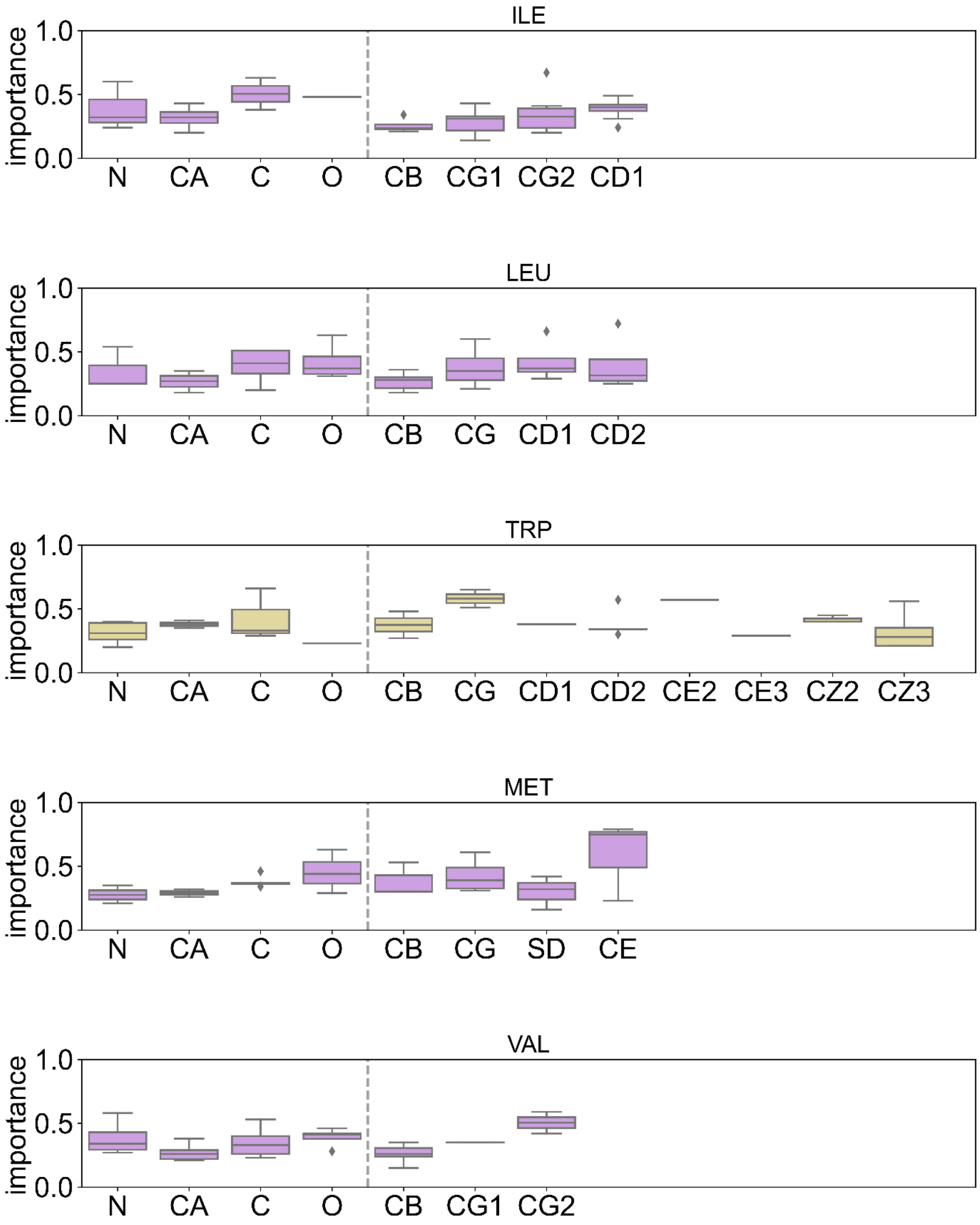
ILE
importance
1.0
0.5
0.0
N
CA
C
O
CB
CG1
CG2
CD1
LEU
importance
N
CA
C
O
CB
CG
CD1
CD2
TRP
importance
N
CA
C
O
CB
CG
CD1
CD2
CE2
CE3
CZ2
CZ3
MET
importance
N
CA
C
O
CB
CG
SD
CE
VAL
importance
N
CA
C
O
CB
CG1
CG2

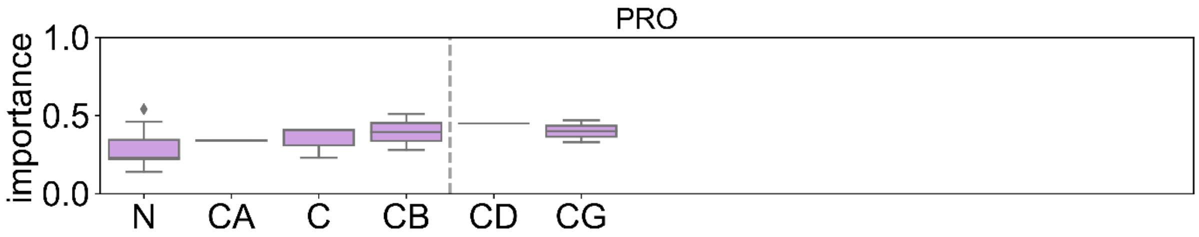
PRO
importance
1.0
0.5
0.0
N
CA
C
CB
CD
CG

**Supporting Figure S17.** Importance for all non-catalytic and non-binding atoms.

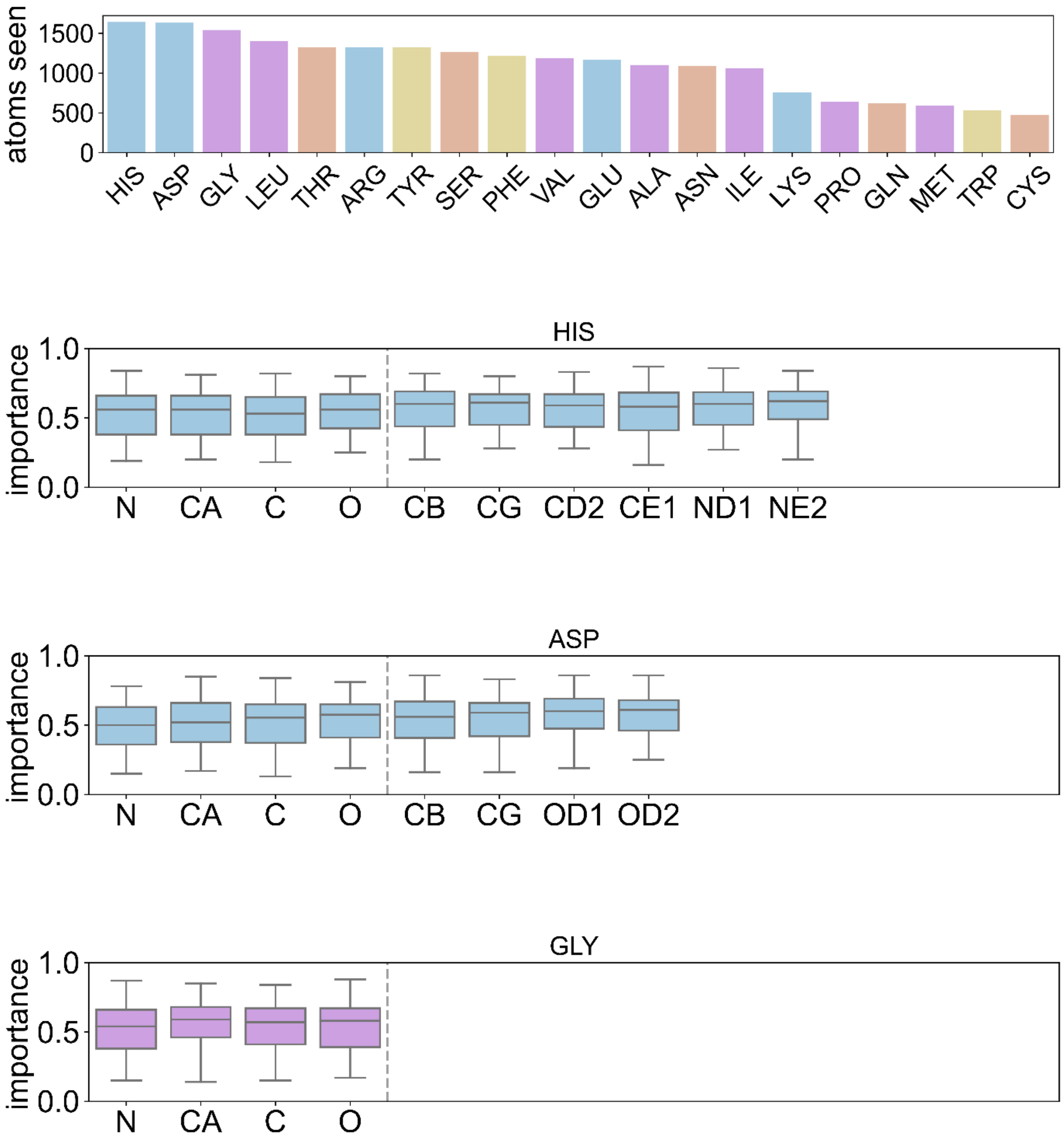

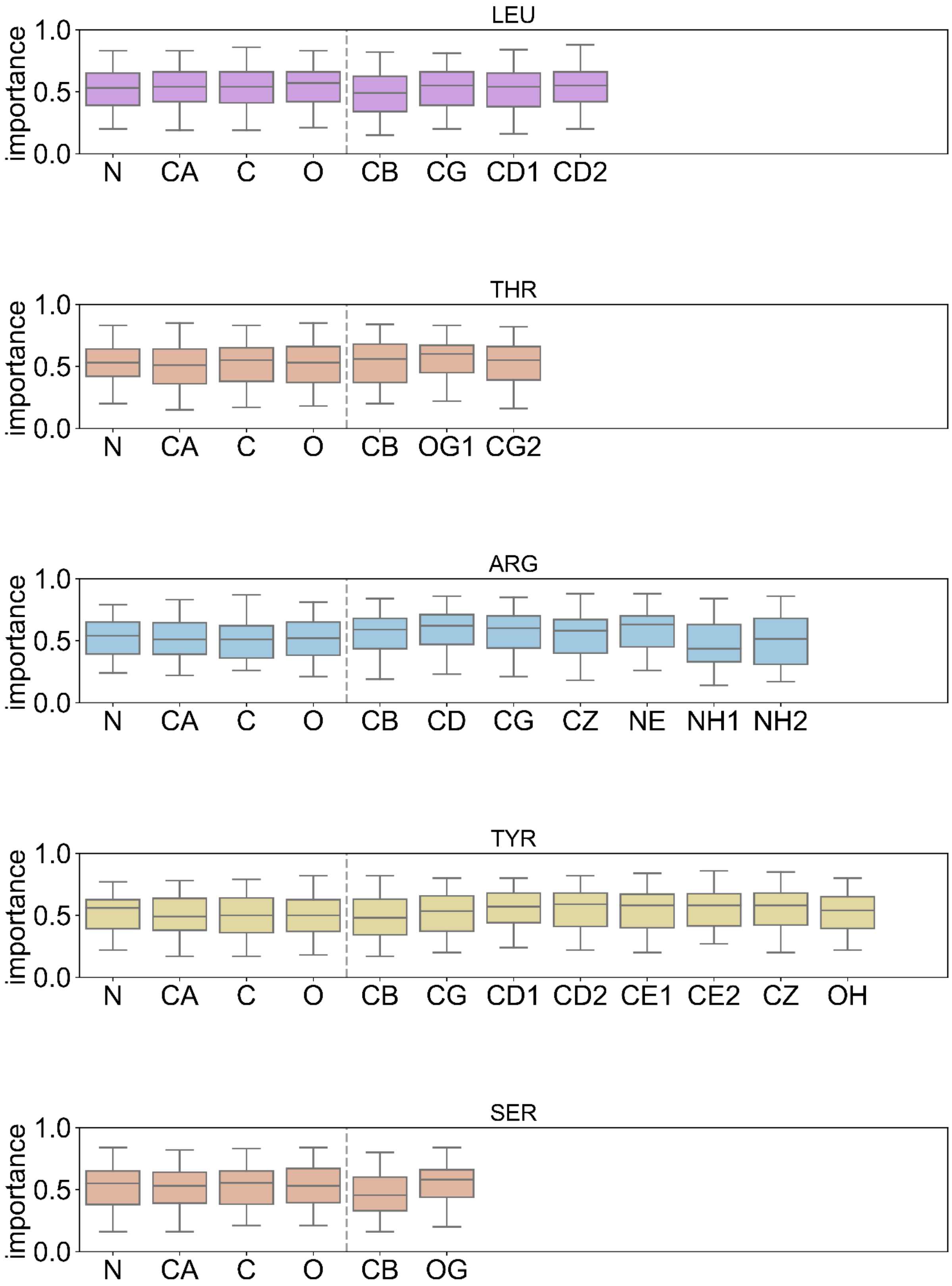
LEU
THR
ARG
TYR
SER
importance
1.0
0.5
0.0
N
CA
C
O
CB
CG
CD1
CD2
OG1
CG2
CD
CZ
NE
NH1
NH2
CE1
CE2
OH
OG

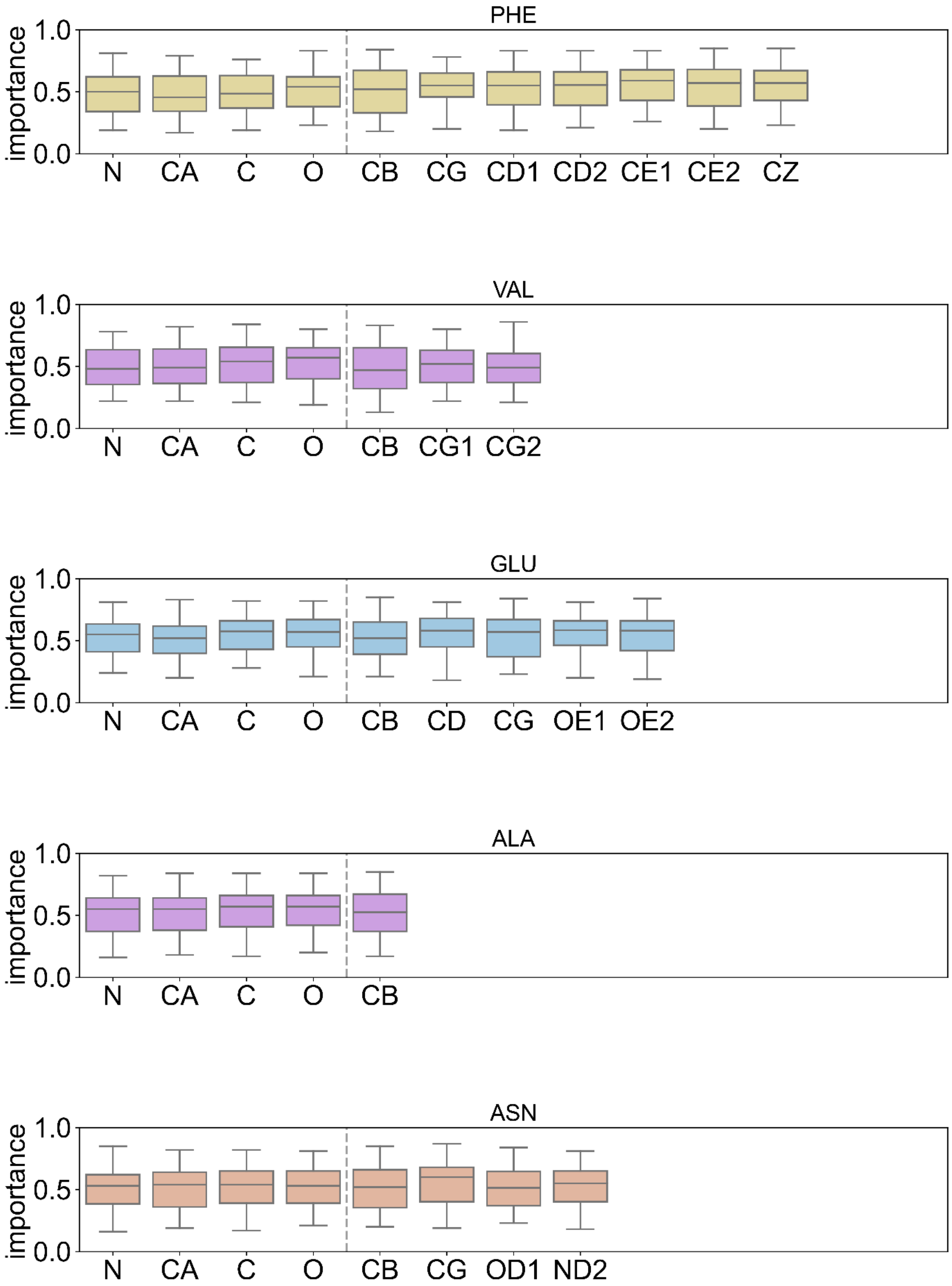
PHE
importance
1.0
0.5
0.0
N CA C O CB CG CD1 CD2 CE1 CE2 CZ
VAL
importance
1.0
0.5
0.0
N CA C O CB CG1 CG2
GLU
importance
1.0
0.5
0.0
N CA C O CB CD CG OE1 OE2
ALA
importance
1.0
0.5
0.0
N CA C O CB
ASN
importance
1.0
0.5
0.0
N CA C O CB CG OD1 ND2

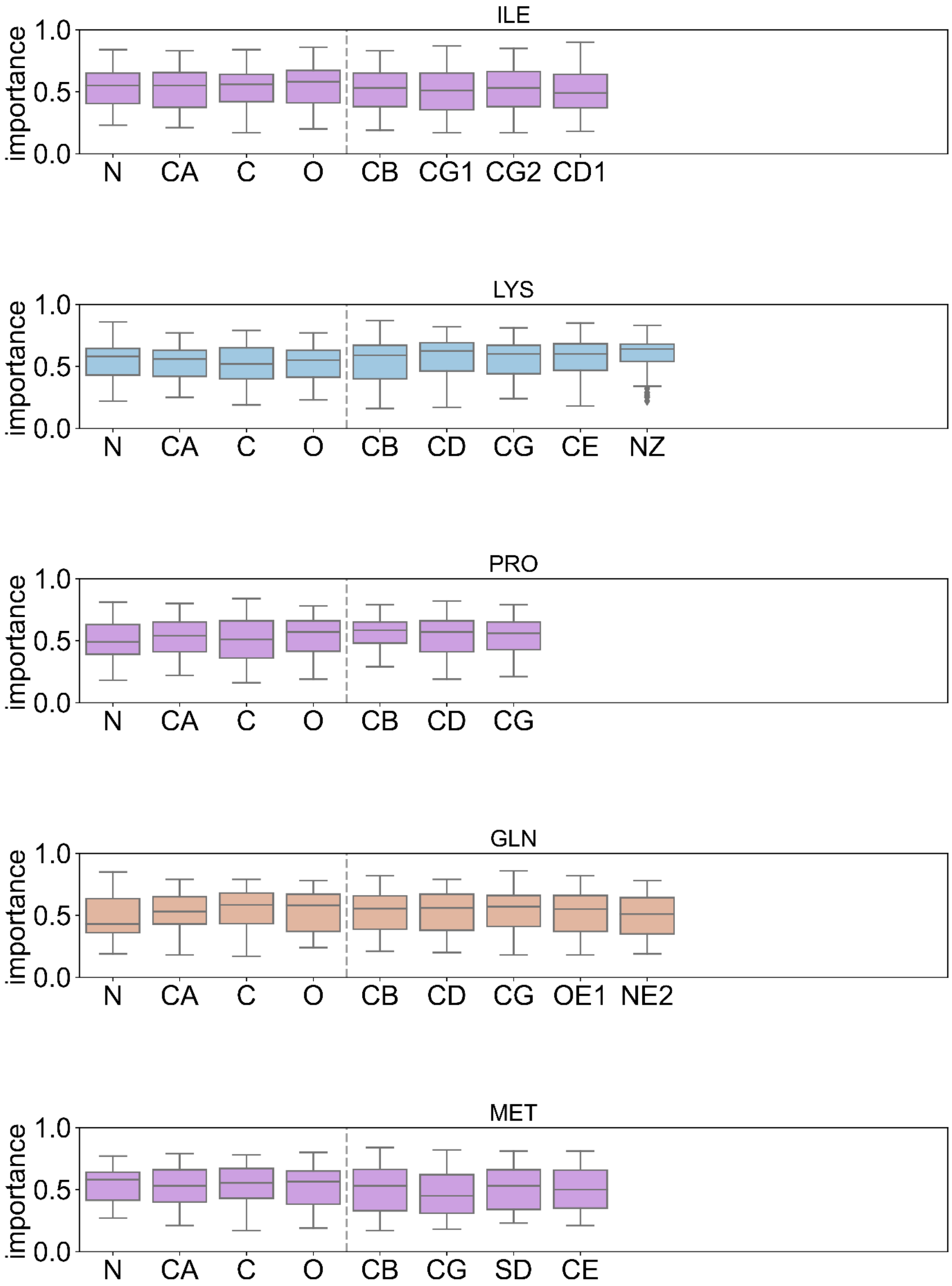

ILE
importance
1.0
0.5
0.0
N
CA
C
O
CB
CG1
CG2
CD1
LYS
importance
1.0
0.5
0.0
N
CA
C
O
CB
CD
CG
CE
NZ
PRO
importance
1.0
0.5
0.0
N
CA
C
O
CB
CD
CG
GLN
importance
1.0
0.5
0.0
N
CA
C
O
CB
CD
CG
OE1
NE2
MET
importance
1.0
0.5
0.0
N
CA
C
O
CB
CG
SD
CE

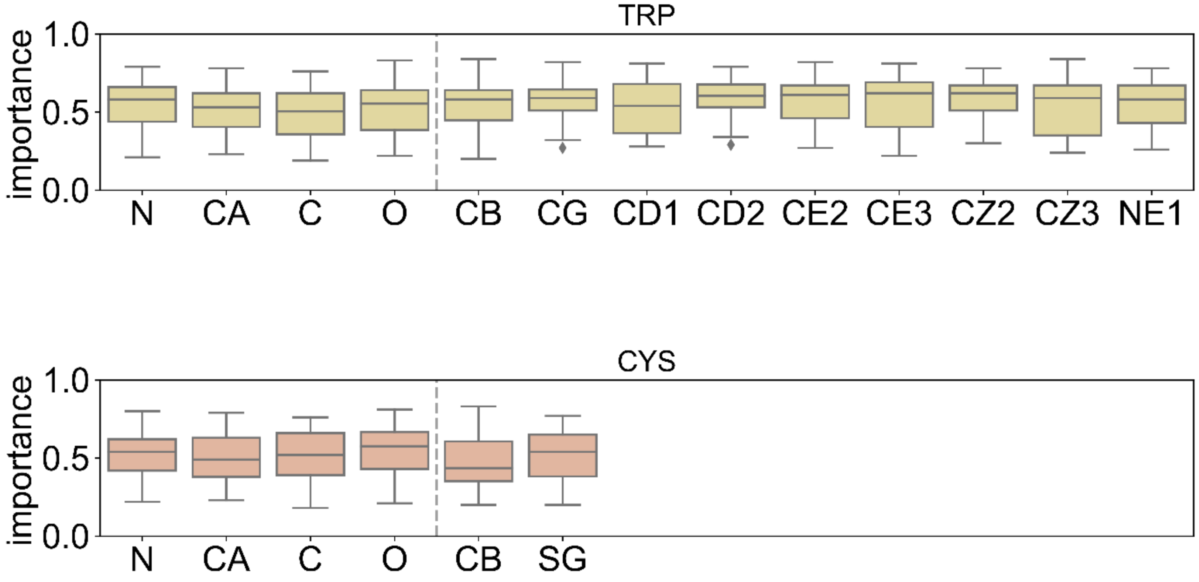
TRP
importance
1.0
0.5
0.0
N
CA
C
O
CB
CG
CD1
CD2
CE2
CE3
CZ2
CZ3
NE1
CYS
importance
1.0
0.5
0.0
N
CA
C
O
CB
SG
